# Efficacy evaluation of glasedgib Sonic Hedgehog pathway inhibition with or without inotuzumab in B-ALL cells using a new co-culturing system model and a validated chemosensitivity assay

**DOI:** 10.64898/2026.05.07.723573

**Authors:** David W. Woolston, Michael Churchill, Carla Grandori, Anjali Advani, Cecilia C.S. Yeung

**Affiliations:** Translational Science and Therapeutics Division, Fred Hutchinson Cancer Center, Seattle, WA, USA; SEngine Precision Medicine, Seattle, WA, USA; Department of Hematology/Oncology, Cleveland Clinic Taussig Cancer Institute, Cleveland, OH, USA; Department of Laboratory Medicine and Pathology, University of Washington Medical Center, Seattle, WA, USA

**Keywords:** Stromal cells, Glasdegib, Sonic Hedgehog pathway, Inotuzumab ozogamicin, *GLI1*, *GLI3*, *PTCH1*, *SMO*

## Abstract

**Purpose:** Glasdegib is a Sonic Hedgehog (SHH) pathway inhibitor used for treating newly diagnosed acute myeloid leukemia in elders or patients unfit for intensive chemotherapy. This study sought to demonstrate growth inhibition and increased apoptosis of B-cell acute lymphoblastic leukemia (B-ALL) in vitro under glasdegib, alone and combined with inotuzumab, using a novel co-culture system and validated chemosensitivity testing model to determine whether glasdegib with and without inotuzumab may represent a promising treatment strategy in B-ALL.

**Methods:** Seven blood and marrow samples from B-ALL patients were co-cultured with HS-5 stromal cells in a co-culturing system designed to mimic the tumor microenvironment to maintain B-ALL cell viability for chemosensitivity testing under glasdegib and inotuzumab.

**Results:** Co-culturing improved B-ALL viability from four to nine days. Dosage-dependent responses to glasdegib were consistent among B-ALL samples on day four based on culture viability, and varied based on expressions of SSH genes GLI1, GLI3, SMO, and PTCH1. Combination with inotuzumab had varied effects on treatment response.

**Conclusion:** Co-culturing B-ALL cells with HS-5 stromal cells improves B-ALL growth and viability. Glasdegib with and without inotuzumab treatments impact the viability of co-cultured B-ALL cells by day four. SHH gene expressions suggest different B-ALL patients may be sensitive or resistant to glasdegib and inotuzumab.

## INTRODUCTION

Although 80-90% of children with acute lymphoblastic leukemia (ALL) are cured (1), more than half of adults with ALL will relapse (2, 3). At the time of relapse, the only known cures are chimeric antigen receptor (CAR-) T cell therapy or an allogeneic hematopoietic stem cell transplant (AHSCT) (4). However, patients typically need to achieve remission before proceeding with these approaches. Both inotuzumab ozogamicin (inotuzumab) (an anti-CD22 antibody drug conjugate) and blinatumomab (an anti-CD19 bispecific antibody) have demonstrated superior rates of response as compared to standard of care chemotherapy in relapsed/refractory precursor (pre)-B-ALL (5, 6). Most B-ALLs express CD22, making inotuzumab an attractive treatment for patients with relapsed/refractory pre-B-ALL. Treatment with inotuzumab has been FDA approved in this setting. Despite high response rates with inotuzumab, the relapse-free survival and overall survival of adult patients remain poor, and further progress is needed to improve outcomes.

Glasdegib is an orally bioavailable small molecular inhibitor of the Sonic Hedgehog (SHH) inhibitor signaling pathway (7). The SHH pathway receptor, patched homologue 1 (PTCH1), controls smoothened (SMO), which is a key activator in the pathway (8). SMO activation results in activation of GLI-dependent transcription factors which can cause proliferation, survival, and differentiation of cells. Inappropriate SMO signaling is implicated in various human malignancies (8), and the SHH pathway plays a key role in proliferation and differentiation of hematopoietic stem cells (8). This pathway has been observed in hematologic malignancies including B-cell lymphomas, multiple myeloma, and leukemia (9). Sonic Hedgehog signaling in the bone marrow stroma is essential for the survival of cancerous B cells through the upregulation of the anti-apoptotic factor, BCL-2 (10, 11). Previous work has demonstrated the importance of the SHH pathway in pre-B ALL (12). Expression of the various pathway components is common in human B-ALL cell lines and clinical samples: 95% of primary B-ALL cell lines expressed PTCH1, GLI1, and SMO (13). In addition, inhibition of the SHH pathway with the SMO inhibitors, cyclopamine or IPI-926, significantly reduced the long-term self-renewal potential in B-ALL leukemic stem cells (13). We hypothesize that treatment with a combination of glasdegib with inotuzumab could potentially demonstrate more effective inhibition of growth and increased apoptosis of cancer cells in B-ALL patients. If this is true, the combination may represent a promsing treatment strategy for relapsed/refractory B-ALL.

In this study, SEngine Precision Medicine used their CLIA-Certified P.A.R.I.S.™ Chemosensitivity Assay, and we added new modifications with a co-culturing system to enabling sustained an *in vitro* growth and viability of B-ALL cells, to evaluate efficacy of SHH inhibition with glasdegib and in combination with inotuzumab in B-ALL patients with high leukemic tumor burden. We then validated a chemosensitivity testing model in a CLIA-certified laboratory to evaluate four treatment schemas.

## MATERIALS AND METHODS

### Samples and materials

B-ALL patient samples obtained from the Fred Hutch/University of Washington Hematopoietic Diseases Repository (IR file #1690) were used. The samples contain fresh collected cryopreserved peripheral bloods (PB) or bone marrow aspirates (BMA) with leukemia burden above 40% blasts. HS-5 stromal cells (ATCC CRL-11882) were used to support B-ALL cell culture in vitro. The study was conducted in accordance with the Declaration of Helsinki and was approved by the Institutional Review Board of Fred Hutchinson Cancer Research Center. Written/signed Fred Hutch IRB-approved consent documents were obtained for all patients in this study.

### Co-culturing assays

To evaluate the ability of human stromal cells in combination with a cytokine cocktail to sustain viability and growth of B-ALL cells. Primary B-ALL blasts were collected either by allowing brief recovery from cryopreserved samples. The leukemic cells were then cultured as described below. The co-culture method is based on RPMI medium with added primocin at 100 μg/mL, FBS at 20% with additional supplements (B27 1X, SCF 50 ng/mL, IL-4 20 ng/mL, CD40 10 ng/mL). Culturing conditions were 37°C with 5% CO_2_. Patients’ cells were cultured *in vitro* with support of a feeder cell layer HS-5 stromal cells (ATCC CRL-11882). HS-5 cells were fluorescently labeled with a blue fluorescent protein to distinguish them from B-ALL cells, as primary cells are difficult to transduce and visually separate from fibroblasts. Biotium NucView® 488 (Biotium, Inc.; Fremont, CA; Cat No. 30029) is a DNA dye that is sequestered in the cytoplasm until caspase 3 is activated. The peptide sequence DEVD is cleaved, allowing the dye to enter the nucleus and bind to DNA. This allows quantification of caspase 3 activation, which indicates apoptosis. The caspase 3 quantification can help determine what percentage of cells are undergoing apoptosis within the blood or bone marrow population, while excluding the HS-5 signal (**Figure 1**). This co-culture system is used to determine the proportion of B-ALL cells undergoing apoptosis upon addition of glasdegib alone and in combination with inotuzumab.

**Figure 1.**
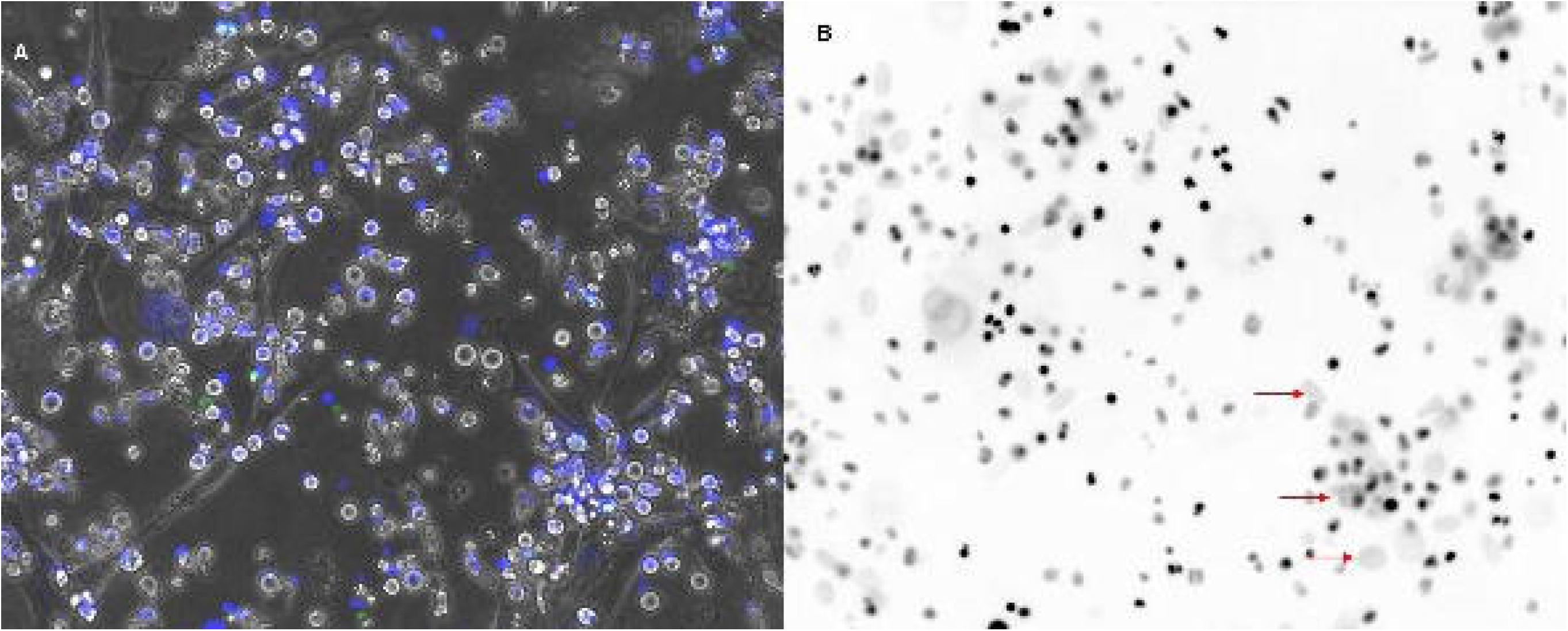
Primary bone marrow cells co-cultured with HS-5 marrow stromal cells. (A) Visualization of the culture in 1 μM Hoechst 33258 (blue) and 0.5 μM NucView (green, indicating activated caspase 3 in cells undergoing apoptosis. (B) Image analysis inverting the blue channel from part A to differentiate HS-5 cells (large diffuse; red arrow) from bone marrow cells (small, roundish, and intense).

### Chemosensitivity assay

The co-culture system was adapted to a chemosensitivity assay under the following conditions. Patient-derived mononuclear cells were thawed (frozen samples) or biopsied (fresh samples) and allowed to recover in co-culture for 48 hours prior to drug treatment under stromal support from a direct contact monolayer of HS-5 feeder cells to assure consistent input viability across all samples. Primary B-ALL cells were observed to be best supported by a confluent monolayer of stromal support cells, but sub-confluent stromal support was sufficient to maintain viability for the assay duration and proved advantageous during mechanical separation.

Glasdegib (DMSO, 10 mM) and inotuzumab (water, 1 μg/mL) were added to collagen-coated 96-well microtiter assay plates using nano volume acoustic droplet ejection (Labcyte® Echo® 555 Liquid Handler; Labcyte Inc.; San Jose, CA) sufficient to achieve final working concentrations and covered with 20 μL RPMI to prevent evaporation and/or hydration. All assay wells were backfilled to a consistent 0.2% DMSO final concentration irrespective of treatment condition.

Viable hematopoietic blasts were isolated with mechanical separation, and their purity was confirmed by light microscopy. Four thousand such mononuclear cells were plated concurrently with 2000 virgin HS-5 stromal cells per well to the assay plate containing drugs to achieve final working volume (100 μL).

Culturing conditions were 37°C with 5% CO_2_. After four days, hematopoietic cells and stroma were mechanically separated, and purity confirmed by light microscopy. Viability relative to vehicle-only control was assessed by ATP quantification using CellTiter-Glo® Luminescent Cell Viability Assay (Promega; Fitchburg, WI; Cat No. G7570), an *in vitro* chemosensitivity testing by a CLIA-certified laboratory was used to determine growth and apoptosis kinetics.

### Data analysis of co-culture viability

Relative viability for each condition was calculated by normalizing total ATP-mediated luminescence to that of cells treated with vehicle (0.2% DMSO) only. Where co-cultured B-ALL is referenced, B-ALL cells were mechanically separated from the stromal monolayer following glasdegib and/or inotuzumab-ozogomycin exposure and transferred to a secondary assay plate for quantification; viability of isolated B-ALL cells was determined as above. In either case, technical triplicates were collected for each dose and growth condition, vehicle-only conditions were replicated 10x for each growth condition. Statistical significance was assessed using Student’s *t*-test.

### Quantitative polymerase chain reaction (qPCR)

Sonic Hedgehog gene expression was tested by quantitative polymerase chain reaction (qPCR) on four genes: *GLI1, GLI3, SMO*, and *PTCH1*. RNA was extracted from cells and cDNA was synthesized with Maxima H minus master mix (ThermoFisher Scientific) using 16 µL of RNA and 4 µL of master mix then incubated at 25°C for 10 min, 50°C for 30 min, and 85°C for 5 min. PCR was carried out using PowerUp™ SYBR™ Green Master Mix (Applied Bioystems; Waltham, MA; Cat No. A25742) on a QuantStudio™ 5 Real-Time PCR System (384-well) thermal cycler (Applied Bioystems; Waltham, MA; Cat No. A28140) with an initial incubation at 50°C for 2 min, 95°C for 10 min, then followed by 40 cycles of 95°C melt for 15 sec, extension at 60°C for 1 min, and ending with a melt curve step consisting of a 95°C melt for 15 sec, incubation at 60°C, then gradual temperature increase to 95°C at 0.075°C/sec. Data was analyzed with the QuantStudio™ Design and Analysis Desktop Software version 1.5.0. SHH gene expressions were quantified using 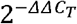 method (14) with *GAPDH* as the reference gene (i.e., internal control) and DMSO-treated control samples as the calibrator against which fold change in gene expression is measured:

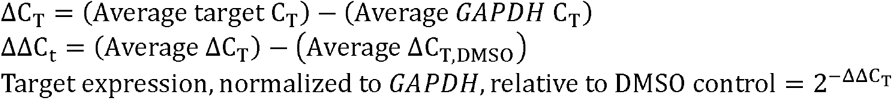

Per conventional qPCR practice, *C*_*T*_ denotes the “threshold cycle”.

Primer sequences follow from 5’ to 3’ and are presented in **Table 1**. Primers for *GLI3* and *SMO* primers were designed with PrimerBank (15). All other primers were designed by hand based on the sequence of the gene transcripts.

**Table 1.**
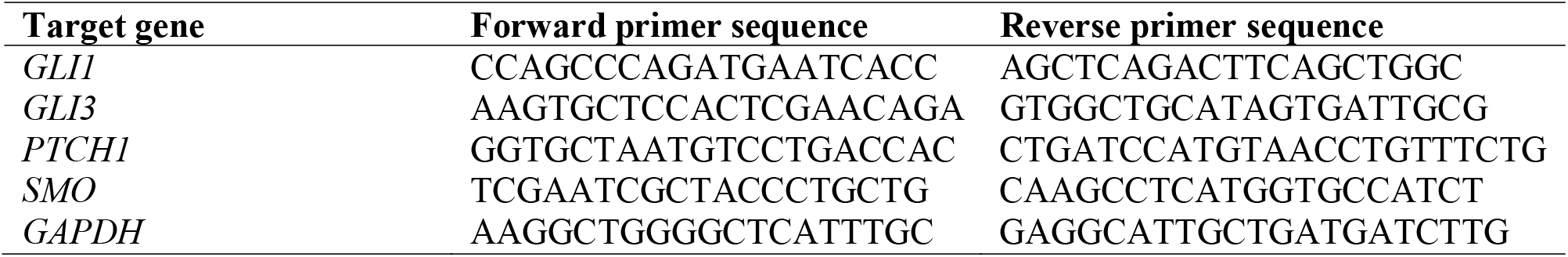
Primer sequences for *GLI1, GLI3, PTCH1, SMO*, and *GAPDH*.

**Table 2.** Patient demographics and sample information for this study’s cohort of seven patients with B-ALL.

| Patient # | Sample type | Blast % | Age at sample collection | Gender | Diagnosis |
| --- | --- | --- | --- | --- | --- |
| 1 | BMA | 96 | 18 | F | B-ALL |
| 2 | PB | 86 | 23 | M | B-ALL |
| 3 | PB | 65.4 | 45 | F | B-ALL |
| 4 | PB | 96 | 30 | F | B-ALL |
| 5 | PB | 42 | 76 | M | B-ALL with secondary MDS |
| 6 | BMA | 63 | 36 | F | B-ALL |
| 7 | BMA | 80 | 26 | M | B-ALL |
| 8 | Stroma | N/A | N/A | N/A | N/A |

M = Male; F = Female; B-ALL = B-Cell Acute Lymphoblastic Leukemia; BMA = Bone Marrow Aspirate; PB = Peripheral Blood

## RESULTS

### Patient and sample cohort

The B-ALL patient samples were derived from four females and three males whose ages ranged from 18– 76 years old. The samples’ blast % ranged from 42–96%. Blasts were isolated from peripheral blood in four samples and bone marrow aspirate in three samples.

### Co-culture of primary B-ALL cells on the SEngine assay and chemosensitivity data

A stromal lawn of HS-5 cells was grown on culturing plates surfaces for two days. Primary B-ALL blasts were recovered from cryostorage and briefly allowed to recover in co-culture with HS-5 stromal support. Experiments showed that Hoechst 33258 (Invitrogen; Thermo Fisher Scientific; Waltham, MA; Catalog No. H3569)—a DNA-specific fluorescent stain—at 1 µM was tolerated by cells for up to eight days in co-culture and allowed for discrimination of HS-5 nuclei from primary B-ALL blasts which generated the workflow schema for our chemosensitivity assay (**Figure 2A**). Viable blasts were plated for chemosensitivity testing in co-culture and exposed to treatments for four days. After four days, blasts were isolated and assessed for viability using CellTiter-Glo.

**Figure 2.**
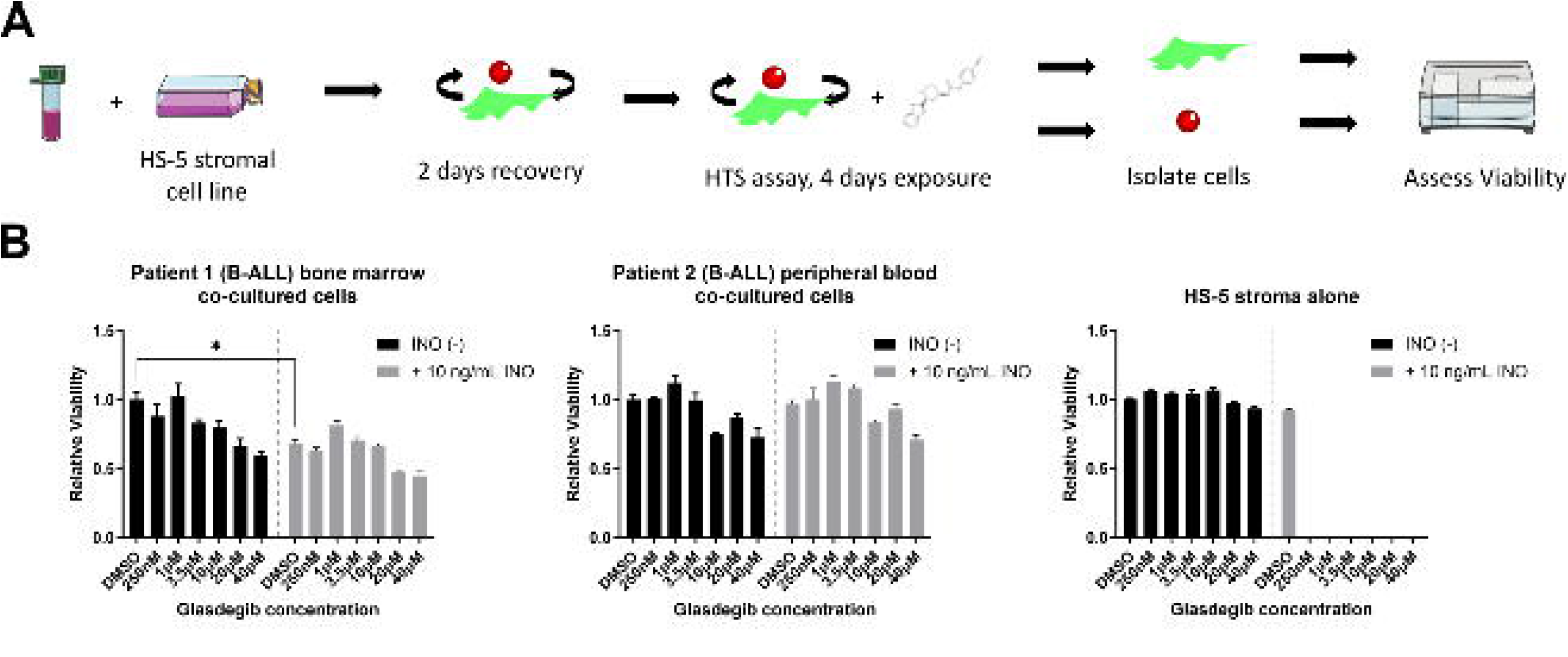
Effect of co-treatment of B-ALL cells with glasdegib and inotuzumab. (A) Experimental design: primary B-ALL blasts were recovered from cryostorage and briefly allowed to recover in co-culture with HS-5 stromal support. Viable blasts were plated for chemosensitivity testing in co-culture and exposed to treatments for four days. Hematopoietic cells were mechanically isolated and assessed for viability using CellTiter-Glo. (B) Relative viability: dose-dependent responses to glasdegib are in seen within the hematopoietic compartment of co-cultured cells (Patients 1 and 2) but not in HS-5 stromal cells alone. Unique patients have differential responses to inotuzumab alone, but co-treatment sensitizes blasts to the effect of glasdegib.

Represented in **Figure 2B** are one bone marrow specimen (patient 1 (**Supplemental Table 1**)) and one peripheral blood specimen (patient 2 (**Supplemental Table 2**)). Individual patient samples responded uniquely to the various treatment schemas. For example, patients 1 and 2 show differential responses to inotuzumab alone, but co-treatment sensitizes blasts to the effect of glasdegib in both samples. This differential sensitivity to inotuzumab did not correlate to peripheral blood specimens throughout the cohort and was infrequently observed. Responses to glasdegib were restricted to the hematopoietic compartment of co-cultured cells; HS-5 stromal cells alone showed negligible sensitivity (**Figure 2B, Supplemental Table 1**).

A consistent dose-dependent response to glasdegib was observed across the cohort (**Figure 3**). While all primary B-ALL blasts demonstrated a consistent sensitivity to glasdegib, only a few samples responded to inotuzumab alone (this was expected due to using a low inotuzumab concentration), and none showed a clear synergistic enhancement to co-treatment of the two agents (**Figure 3B**).

**Figure 3.**
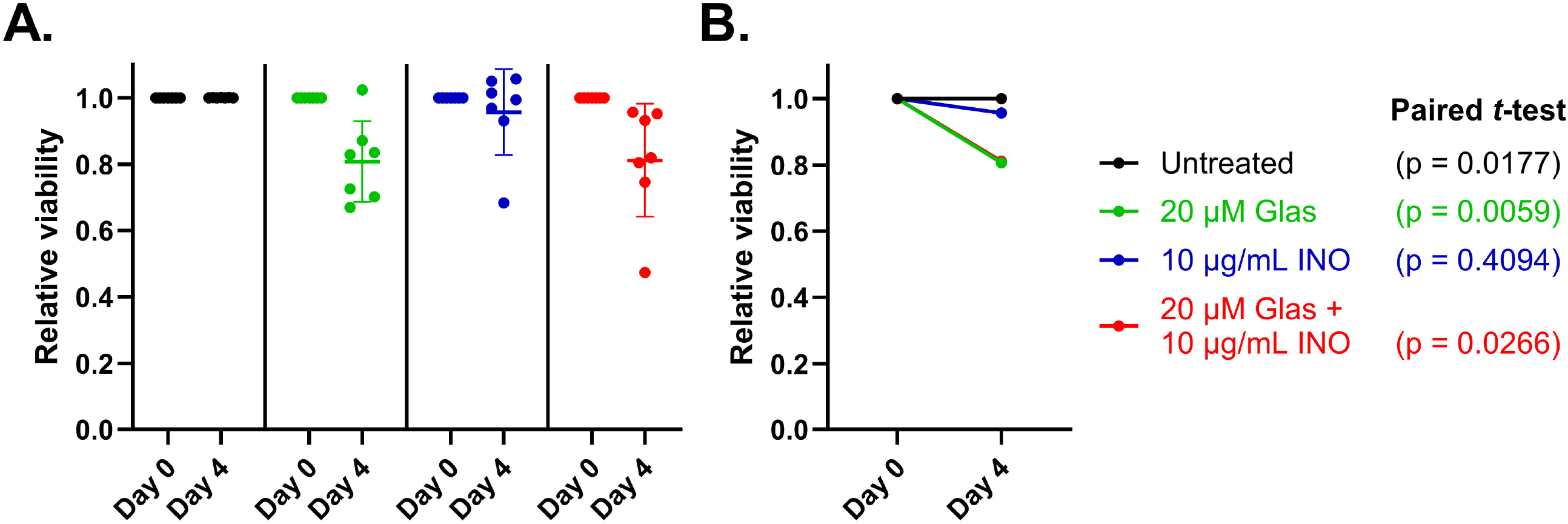
Seven primary B-ALL samples assessed for viability using CellTiter-Glo at day 0 and at day 4. The x-axes represent the post-culture timepoint. The y-axes represent B-ALL samples’ viability relative to and normalized against a control sample. Four conditions are tested, each represented by different colors in the scatter plots. Significant loss of viability was observed after four days under treatment with glasdegib, inotuzumab, and combination glasdegib/inotuzumab. (A) Culture viability under four treatment conditions with each point representing one of seven primary B-ALL samples at day 0 and day 4. Each point represents the meaning of triplicate viability fractions for each patient. Although statistically significant separation was observed for the untreated control samples, it should be noted the untreated samples’ Δ average relative viability was +0.00005711, compared with -0.1923 for glasdegib-treated samples and -0.1881 for glasdegib + inotuzumab-treated samples (in both cases, a < -3000-fold difference). (B) Mean viability under each treatment condition at day 0 and day 4. Note there is a commensurate decrease in viability under glas 20 μM (green) and glas 20 μM + INO 10 μg/mL (red, hidden under the green line) treatments.

### Sonic Hedgehog gene expression

The analysis of SHH gene expression was challenged by the initial viability reduction in the B-ALL blasts in the co-culturing system. Of the remaining five samples, four provided enough RNA for expression analysis (patients 3, 5, 6, and 8).

Figure 4. visualizes the primary B-ALL specimens’ SHH gene expressions at day 4 post co-culture. The day 4 timepoint was studied to specifically assess expression when viability was reduced. Patient 3 (**Figure 4A**) presents a patient sample which did not respond well to any treatment conditions and appears to be resistant to both glasdegib 20 μm (glas 20) and glasdegib 20 μM + inotuzumab (glas 20 + INO); in fact, there appeared to be overexpression of *GLI3* and *SMO* in response to treatment with glas 20 and glas 20 + INO. Patient 5 (**Figure 4B**) showed a small decrease in *GLI1* with only glas 20 + INO. Patient 6 (**Figure 4C**) showed decreased *GLI1* and *PTCH1* under both glas 20 and glas 20 + INO and may indicate a synergistic effect between the two drugs given that glas 20 + INO demonstrated complete loss of expression in this patient’s sample. Patient 7 (**Figure 4D**) demonstrated a minor decrease with glas 20 but appears to have synergistic downregulation of *FLI1* under combination glas 20 + INO. Although statistical analysis could not be performed given small sample sizes due to limited material, we observed three out of four primary B-ALL specimens which underwent this chemosensitivity assay demonstrate variation in SHH signaling pathway gene expression of *GLI1, GLI3, SMO*, or *PTCH1* after treatment with glasdegib alone or with inotuzumab.

## DISCUSSION

We have demonstrated a novel strategy to perform chemosensitivity testing on primary patient-derived B-ALL cells with a novel co-culturing assay capable of extending the life of primary B-ALL cells for over eight days. To this end, after cryo-recovery and brief expansion with full stromal support (“priming”), we isolated primary patient B-ALL samples and transitioned to co-culturing methods with recombinant stromal cell and cytokine support for a drug challenge trial with a novel treatment combination of a SHH inhibitor (glasdegib) with an anti-CD22 agonist (inotuzimab). Initial challenges of maintaining viability in monoculture of B-ALL were improved when primary B-ALL cells are co-cultured on a monolayer of HS-5 stromal cells in combination with a cytokine cocktail. Using this system, we observed reduced viability in primary B-ALL cells treated with higher concentrations of glasdegib, with increased synergistic killing with combination inotuzumab in a subset of the primary patient B-ALL samples.

**Figure 4.**
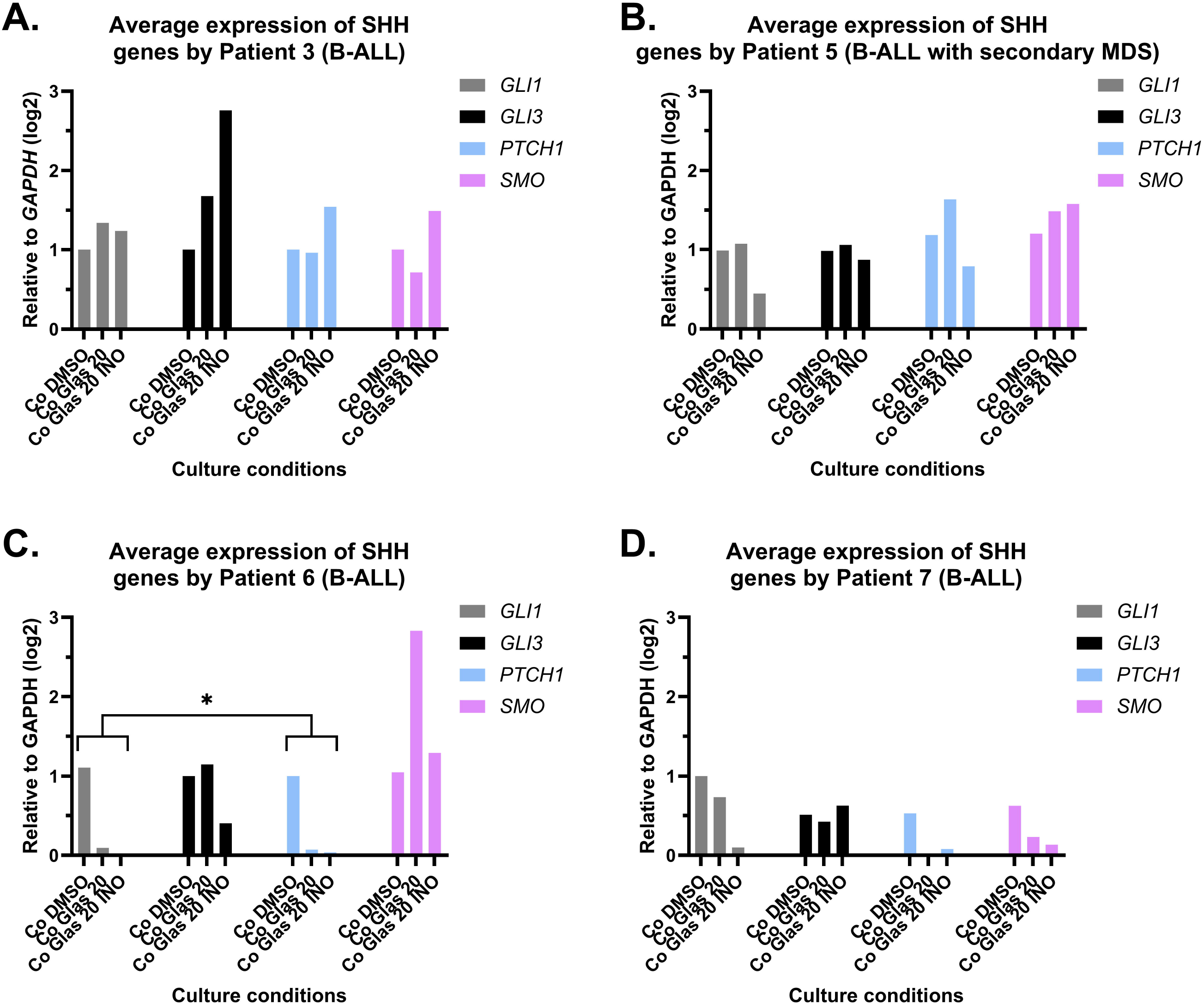
The SHH gene expressions by four primary B-ALL patient samples in different culture conditions. Each panel represents a different patient sample, and each color corresponds to the SHH gene indicated in the legend. The x-axes represent the treatment conditions: negative control (Co DMSO), 20 μM glasdegib (Co Glas 20), and 20 μM glasdegib + 10 ng/mL inotuzumab (Co Glas 20 INO). The y-axes represent the expression of *GLI1, GLI3, SMO*, and *PTCH1* after four days of HS-5 stromal cell co-culture. All expression values were represented by 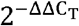 method with *GAPDH* as the internal control and treatment with DMSO as the fold-change calibrator.

Primary hematopoietic cells, even those malignantly transformed, maintain obligatory dependence on their microenvironment for long term ex-vivo culture (16). However, primary leukemias can typically be supported for short periods of time with exogenous cytokine support alone without any insult to their renewal capacity or tumorigenicity (17). The co-culture method described here is the first of its kind enabling sustained primary B-ALL cells to extend viability to eight days which allowed for chemosensitivity testing in B-ALL when adapted to SEngine’s culturing platform allowing for a highly sensitive detection of cell death kinetics. This capacity can be exploited to efficiently measure the effects of experimental conditions such as drug challenges to leukemia cells alone using terminal readouts that do not segregate distinct cell populations such as Cell-Titer-GLO. The CellTiter-Glo® Luminescent Cell Viability Assay is a homogeneous method of determining the number of viable cells in culture based on quantitation of the ATP present and involves adding the single reagent directly to cultured cells with medium, washing, and removing medium. The luminescent signal generated is proportional to the amount of ATP present, which is directly proportional to the number of cells present in culture. We used the level of ATP detected by the CellTiter GLO assay to determine the proportion of B-ALL cells undergoing apoptosis.

Unique challenges in the development of chemosensitivity testing for combination glasdegib with inotuzumab included the inability to use monoculture methods and the need for a stromal component to demonstrate the impact of glasdegib on SHH inhibition. While specialized cytokine and supplement cocktails sufficiently supported the leukemic cells in monoculture for a brief assay duration, due to the mechanisms of action of these two drugs, observing activity the assay itself was less straightforward than initially expected. The combination of CD22+ B-ALL hematopoietic cells with associated stroma must be co-cultured to realize the effect of glasdegib-mediated inhibition of paracrine SHH signaling. This necessitated the development of an extended culturing assay described in this work to elucidate the effects of glasdegib on B-ALL viability in-vitro. In this study, we used a lower inotuzumab concentration to show more effect for SHH inhibition and therefore could not demonstrate significant synergistic effects in all samples. As such, we can show variance from baseline (DMSO) in the primary B-ALL leukemic cells. We chose to use DMSO as the comparator since this control sample would be treated in the exact same manner in every way except for the addition of drug therapy.

## CONCLUSION

In summary, this early proof of concept study observed viability reduction and gene expression differences that correspond to treatment with 20 μM glasdegib in a small pilot cohort of B-ALL patient samples. Our data support that treatment with glasdegib and combination glasdegib + inotuzumab impacts viability of primary B-ALL cells in a co-culturing chemosensitivity assay by day four. Possible clinical implementation of these findings could include a workflow where gene expression analysis is performed on patient samples with B-ALL. Given specific gene expression profiles patients, can be stratified into those who may response well to glasdegib and inotuzumab, e.g., specifically identifying patients who are sensitive versus patients who may be resistant.

## ACKNOWLEDGEMENTS

Thanks to the Torok-Storb Lab for assisting in developing the new co-culturing system and SEngine laboratory in performing CellTiter-Glo and the Chemosensitivity assays. We also appreciate the patients who enrolled in the Fred Hutch Hematologic Malignancy Repository program. We are grateful to have SWOG as our partner and provide funding for this research.

## REFERENCES

1. M. J. Burke, M. Devidas, Z. Chen, W. L. Salzer, E. A. Raetz, K. R. Rabin, N. A. Heerema, A. J. Carroll, J. M. Gastier-Foster, M. J. Borowitz, B. L. Wood, N. J. Winick, W. L. Carroll, S. P. Hunger, M. L. Loh and E. C. Larsen (2022) Outcomes in adolescent and young adult patients (16 to 30 years) compared to younger patients treated for high-risk B-lymphoblastic leukemia: report from Children’s Oncology Group Study AALL0232. Leukemia 36, 648–655.

2. A. Oriol, S. Vives, J. M. Hernandez-Rivas, M. Tormo, I. Heras, C. Rivas, C. Bethencourt, F. Moscardo, J. Bueno, C. Grande, E. del Potro, R. Guardia, S. Brunet, J. Bergua, T. Bernal, M. J. Moreno, C. Calvo, P. Bastida, E. Feliu, J. M. Ribera and G. Programa Espanol de Tratamiento en Hematologia (2010) Outcome after relapse of acute lymphoblastic leukemia in adult patients included in four consecutive risk-adapted trials by the PETHEMA Study Group. Haematologica 95, 589–596.

3. E. Jabbour, S. O’Brien, M. Konopleva and H. Kantarjian (2015) New insights into the pathophysiology and therapy of adult acute lymphoblastic leukemia. Cancer-Am Cancer Soc 121, 2517–2528.

4. T. H. Truong, C. Jinca, G. Mann, S. Arghirescu, J. Buechner, P. Merli and J. A. Whitlock (2021) Allogeneic Hematopoietic Stem Cell Transplantation for Children With Acute Lymphoblastic Leukemia: Shifting Indications in the Era of Immunotherapy. Front Pediatr 9, 782785.

5. H. M. Kantarjian, D. J. DeAngelo, M. Stelljes, G. Martinelli, M. Liedtke, W. Stock, N. Gokbuget, S. O’Brien, K. Wang, T. Wang, M. L. Paccagnella, B. Sleight, E. Vandendries and A. S. Advani (2016) Inotuzumab Ozogamicin versus Standard Therapy for Acute Lymphoblastic Leukemia. N Engl J Med 375, 740–753.

6. H. M. Kantarjian, A. S. Stein, R. C. Bargou, C. Grande Garcia, R. A. Larson, M. Stelljes, N. Gokbuget, G. Zugmaier, J. E. Benjamin, A. Zhang, C. Jia and M. S. Topp (2016) Blinatumomab treatment of older adults with relapsed/refractory B-precursor acute lymphoblastic leukemia: Results from 2 phase 2 studies. Cancer-Am Cancer Soc 122, 2178–2185.

7. N. Fukushima, Y. Minami, S. Kakiuchi, Y. Kuwatsuka, F. Hayakawa, C. Jamieson, H. Kiyoi and T. Naoe (2016) Small-molecule Hedgehog inhibitor attenuates the leukemia-initiation potential of acute myeloid leukemia cells. Cancer Sci 107, 1422–1429.

8. Y. Lim and W. Matsui (2010) Hedgehog signaling in hematopoiesis. Crit Rev Eukaryot Gene Expr 20, 129–139.

9. C. Y. Ok, R. R. Singh and F. Vega (2012) Aberrant activation of the hedgehog signaling pathway in malignant hematological neoplasms. The American journal of pathology 180, 2–11.

10. R. Chaber, L. Fiszer-Maliszewska, D. Noworolska-Sauren, J. Kwasnicka, G. Wrobel and A. Chybicka (2013) The BCL-2 protein in precursor B acute lymphoblastic leukemia in children. J Pediatr Hematol Oncol 35, 180–187.

11. I. Kapoor, J. Bodo, B. T. Hill, E. D. Hsi and A. Almasan (2020) Targeting BCL-2 in B-cell malignancies and overcoming therapeutic resistance. Cell Death Dis 11, 941.

12. Z. Sheybani, S. Rahgozar and E. S. Ghodousi (2019) The Hedgehog signal transducer Smoothened and microRNA-326: pathogenesis and regulation of drug resistance in pediatric B-cell acute lymphoblastic leukemia. Cancer Manag Res 11, 7621–7630.

13. T. L. Lin, Q. H. Wang, P. Brown, C. Peacock, A. A. Merchant, S. Brennan, E. Jones, K. McGovern, D. N. Watkins, K. M. Sakamoto and W. Matsui (2010) Self-renewal of acute lymphocytic leukemia cells is limited by the Hedgehog pathway inhibitors cyclopamine and IPI-926. Plos One 5, e15262.

14. K. J. Livak and T. D. Schmittgen (2001) Analysis of relative gene expression data using real-time quantitative PCR and the 2(-Delta Delta C(T)) Method. Methods 25, 402–408.

15. A. Spandidos, X. Wang, H. Wang and B. Seed (2010) PrimerBank: a resource of human and mouse PCR primer pairs for gene expression detection and quantification. Nucleic Acids Res 38, D792–799.

16. J. Frobel, T. Landspersky, G. Percin, C. Schreck, S. Rahmig, A. Ori, D. Nowak, M. Essers, C. Waskow and R. A. J. Oostendorp (2021) The Hematopoietic Bone Marrow Niche Ecosystem. Front Cell Dev Biol 9, 705410.

17. D. G. J. Cucchi, R. W. J. Groen, J. Janssen and J. Cloos (2020) Ex vivo cultures and drug testing of primary acute myeloid leukemia samples: Current techniques and implications for experimental design and outcome. Drug Resist Updat 53, 100730.

